# Quantitative Analysis of Autophagy in Single Cells: Differential Response to Amino Acid and Glucose Starvation

**DOI:** 10.1101/2023.12.01.569679

**Authors:** Katie R. Martin, Stephanie L. Celano, Ryan D. Sheldon, Russell G. Jones, Jeffrey P. MacKeigan

## Abstract

Autophagy is a highly conserved, intracellular recycling process by which cytoplasmic contents are degraded in the lysosome. This process occurs at a low level constitutively; however, it is induced robustly in response to stressors, in particular, starvation of critical nutrients such as amino acids and glucose. That said, the relative contribution of these inputs is ambiguous and many starvation medias are poorly defined or devoid of multiple nutrients. Here, we sought to generate a quantitative catalog of autophagy across multiple stages and in single, living cells under normal growth conditions as well as in media starved specifically of amino acids or glucose. We found that autophagy is induced by starvation of amino acids, but not glucose, in U2OS cells, and that MTORC1-mediated ULK1 regulation and autophagy are tightly linked to amino acid levels. While autophagy is engaged immediately during amino acid starvation, a heightened response occurs during a period marked by transcriptional upregulation of autophagy genes during sustained starvation. Finally, we demonstrated that cells immediately return to their initial, low-autophagy state when nutrients are restored, highlighting the dynamic relationship between autophagy and environmental conditions. In addition to sharing our findings here, we provide our data as a high-quality resource for others interested in mathematical modeling or otherwise exploring autophagy in individual cells across a population.

## INTRODUCTION

Autophagy is a highly conserved intracellular degradation pathway that recycles cytoplasmic contents through a vesicular trafficking pathway that culminates in the lysosome. Under normal conditions, autophagy serves a housekeeping function in cells by degrading long-lived proteins and clearing damaged organelles. However, during stress, autophagy is activated to liberate internal nutrient pools to support metabolism and circumvent cell death. There is a growing appreciation for the role of this stress response pathway in cancer, where autophagy has been shown to promote the progression of KRAS and BRAF-driven tumors^1–5^ by supporting tumor-intrinsic cancer cell survival, and also, by regulating additional cell types in the tumor microenvironment (i.e., immune cells)^6–9^.

The gatekeeper of autophagy is ULK1, a serine/threonine kinase that triggers the nucleation of an isolation membrane, or phagophore, the earliest autophagic membrane^10–12^. The synthesis of this cup-shaped structure is largely promoted by the class III PI3K (VPS34) and its lipid product, PI(3)P. PI(3)P decorates autophagic membranes and recruits effectors, such as DFCP1 and WIPI family proteins (e.g., WIPI1, WIPI2)^13–15^. VPS34 activity is required for the downstream activation of ATG9, a transmembrane protein that facilitates lipid transport to contribute to the expansion of the phagophore membrane^16–20^. Maturation of the phagophore and eventual closure into a complete autophagosome is executed by two ubiquitin-like conjugation systems. The first system involves covalent binding of ATG12, a ubiquitin-like protein, to ATG5 and subsequent incorporation into a large oligomer with ATG16 at the phagophore^21^. The second system involves the classic autophagosome-marker, LC3 (Atg8 in yeast), which becomes covalently attached to phosphatidylethanolamine on autophagic membranes^22,23^. The location and enabling of LC3 conjugation to the vesicle is controlled by ATG12-5-16, which functions as an E3-like enzyme^24^. The completion of this process is marked by the autophagosome’s direct fusion with a lysosome (generating an autolysosome), or with an endosome destined for the lysosome (generating an amphisome), and the degradation of sequestered cargo.

Autophagy is tightly regulated by environmental conditions, most notably, it is activated by amino acid stress through regulation by mammalian target of rapamycin (MTOR; specifically MTOR complex 1 or MTORC1) (**Fig. 1A**). Under amino acid sufficiency, activated MTORC1 restrains autophagy through inhibitory phosphorylation of the ULK1 complex (at Ser758)^10–12,25,26^. This serves to limit autophagy-mediated catabolism during times of sufficient exogenous nutrient supply. Upon amino acid withdrawl, MTORC1 is repressed, thereby relieving its inhibition of ULK1 and activating autophagy. Autophagy can also be activated by energetic stress through regulation of the energy-sensing AMP-activated protein kinase (AMPK) (**Fig. 1A**). Under a favorable energy status, AMPK, which promotes ULK1 activity through activating phosphorylation of multiple sites, is inhibited, which contributes to autophagy suppression. During energetic stress (i.e., decreased cellular ATP/AMP ratio), AMPK is activated to inhibit MTORC1, while also thought to directly activate ULK1 through phosphorylation^27–29^. Despite this, the precise contribution of glucose starvation to autophagy is not well understood and the dogma that AMPK activation increases ULK1-dependent autophagy during energetic stress has been challenged in recent years^30–33^.

**Figure 1.**
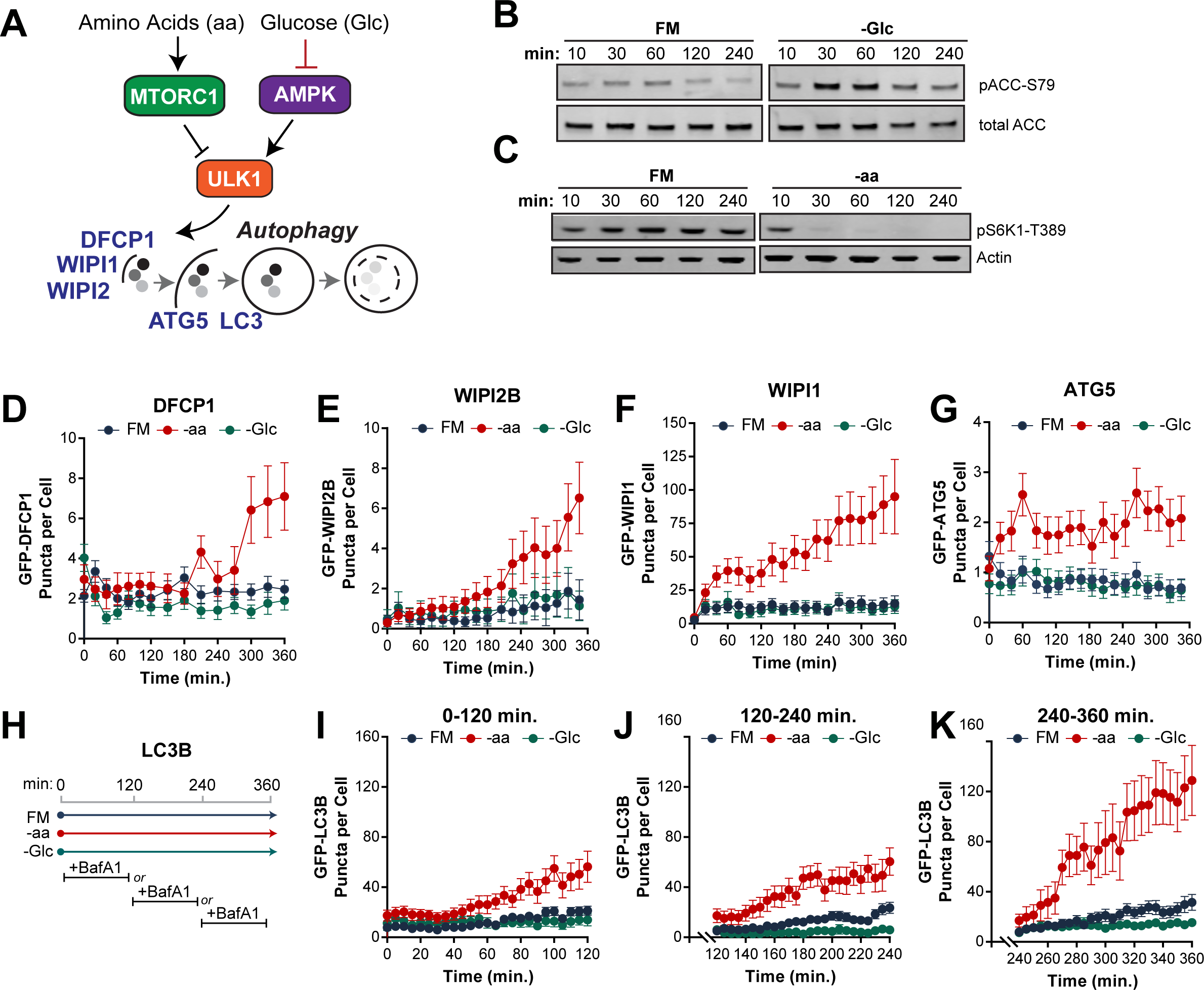
Amino acid starvation, but not glucose starvation, induces autophagy in U2OS cells. (**A**) MTORC1 and AMPK activity is influenced by amino acids and glucose, respectively, and regulate ULK1 activity, which induces autophagy. (B-C) U2OS cells were treated with full media (FM), FM lacking glucose (-Glc), or FM lacking amino acids (-aa) for the indicated times and cell lysates analyzed by immunoblot for ACC, total and phosphorylated at S79 (**B**), and S6K1 phosphorylated at T389 and actin as a loading control (**C**). (D-G) Monoclonal U2OS cell lines expressing DFCP1 (**D**), WIPI2B (**E**), WIPI1 (**F**), or ATG5 (**G**) were treated with FM (blue), -Glc (green), or -aa media (red) for 6 hours and subjected to live-cell fluorescent imaging. GFP-puncta (objects) for each reporter were quantified from single cells. Trajectories include mean objects per cell (symbols); bars represent 95% CI. (H-K) GFP-LC3B objects were quantified from cells treated with FM, -aa, or -Glc in the presence of BafA1 (to prevent lysosome degradation) in 2 hour increments (**H**): 0-120 min (**I**), 120-240 min (**J**), and 240-360 min (**K**). Trajectories include mean objects per cell (symbols); bars represent 95% CI.

To provide clarity on the impact of nutrient stress on autophagy, we used a well-characterized cancer cell line, U2OS, and fluorescent sensors across multiple stages of autophagy to monitor autophagy in response to starvation or amino acids or glucose. For this, we formulated defined medias to precisely limit these nutrients in isolation while controlling all other media components. Using these medias, we found that withdrawal of amino acids, but not glucose, induced autophagy in U2OS cells. Moreover, we confirmed that MTORC1 regulation of ULK1 and the magnitude of autophagy are tightly linked and correlate with amino acid availability. Moreover, we found that while autophagy is activated immediately in response to amino acid deprivation, further upregulation occurs several hours later, concomitant with transcriptional upregulation of autophagy machinery. Finally, we demonstrate that cells are primed to return to a low-autophagy state when nutrients are replenished. In addition to these well-supported conclusions, we provide kinetic, single-cell data from living cells captured across multiple stages of autophagy as a resource.

## RESULTS

### Starvation of amino acids, but not glucose, induces robust autophagy in U2OS cells

To understand the relative contribution of the major physiologic inputs to autophagy, we measured multiple stages of the pathway in U2OS cells cultured in each of three defined medias (see **Supplemental Table 1** for full formulations): *1)* custom full media (“FM”), which contains the concentrations of amino acids, growth factors, glucose, and vitamins found in RPMI-1640; *2)* amino acid starvation media (“-aa”), which is FM without amino acids; and *3)* glucose starvation media (“-Glc”), which is FM without glucose. First, we confirmed that -Glc media increased AMPK activity as measured by phosphorylation of the AMPK substrate, ACC, at Ser79), an effect that was immediate and maintained throughout the entire 6 hour period (**Fig. 1B**). We also established that amino acid starvation inhibited MTORC1, as indicated by a complete loss of phosphorylation of its substrate, S6K1, at T389 within 30 minutes (**Fig. 1C**).

To determine the consequences of these medias on autophagy, we used a panel of monoclonally derived U2OS cell lines expressing EGFP-fusions of key autophagy machinery including DFCP1, WIPI1, WIPI2, ATG5, and LC3B (**Fig. 1A**). Cells were plated in FM and then switched to either FM, -aa, or -Glc before fluorescent imaging live cells over the course of 6 hours. We found that fed cells (cultured in FM) generally expressed few (2.1 +/-1.4) DFCP1-positive puncta, which represent omegasomes, the crescent-shaped structure that supports the forming phagophore, and these were relatively stable through the 6 hour imaging period (**Fig. 1D, blue**). Amino acid starvation (-aa) increased DFCP1-positive puncta, primarily after 3 hours of treatment, peaking at an average of 7.1 puncta per cell by 6 hours (**Fig. 1D, red**). In contrast, -Glc failed to induce DFCP1 puncta formation (**Fig. 1D, green**). WIPI2B-positive puncta were also lowly abundant and responded similarly to -aa, rising from 0.3 to 6.5 puncta per cell on average (>20-fold), most substantially in the final 3 hours of starvation, while failing to respond to -Glc as well (**Fig. 1E**). The second PI(3)P effector we quantified, WIPI1, also responded exclusively to -aa; however, this marker showed an immediate increase in puncta, rising from approximately 5 to 40 puncta per cell in the first hour of -aa, followed by a slow and steady increase through the remainder of the starvation period (**Fig. 1F**). Similar to WIPI1, the first hour of -aa triggered a doubling in the number of ATG5-positive puncta, although these structures were overall very rare (1-2 per cell) (**Fig. 1G**).

As LC3B is known to be conjugated to the growing autophagic membrane and degraded in the lysosome along with cargo, measuring it requires a more complex experimental design. Bafilomycin A1 (BafA1) is a proton pump inhibitor that prevents lysosomal degradation of autophagic vesicles; therefore, the rate at which vesicles accumulate upon BafA1 treatment serves as a proxy for their rate of synthesis^34^. Because the duration of BafA1 treatment must be limited, we spiked in BafA1 for either the first, middle, or final two hours of the 6-hour treatment period and quantified LC3-positive vesicles (**Fig. 1H**). Similar to the other markers, we found that -aa, but not -Glc, caused an increase in LC3-positive vesicles, an effect that was apparent immediately and grew more robust over time (**Fig. 1I**). Taken together, these data strongly suggest that autophagy is upregulated in U2OS cells in response to starvation of amino acids, but not glucose withdrawal.

### MTORC1-mediated autophagy is tuned to amino acid levels

After observing that amino acids are the dominant driver of autophagy in U2OS cells, we wanted to more precisely establish the relationship between amino acid levels and autophagy induction. For this, we titrated amino acids from 100% (the level of each amino acid in the RPMI-1640 formulation and our custom FM) down to 10%, 5%, and 0%, while keeping all other media components constant. At the end of the 6 hour treatment period, we saw that decreasing amino acids led to decreasing levels of pULK1-S758, with 0% aa causing a near complete loss of pULK1-S758, similar to that observed with pS6K1-T389 in Figure 1 (**Fig. 2A**). These data were fit to a sigmoidal dose-response curve, which revealed an EC_50_ of 6.4% aa (+/-1.2% aa) (**Fig. 2B**). We then measured LC3-positive vesicle synthesis by supplementing BafA1 into these medias and imaging them between 4 and 6 hours of treatment. We found that reduction of amino acids in the media increased the rate of LC3 accumulation, with an EC_50_ of approximately 7.6% aa (+/-2.5% aa) (**Fig. 2C-D**). We confirmed that there was a linear negative relationship between pULK1-S758 and LC3 vesicle synthesis, consistent with relief of this phosphorylation site on ULK1 being a critical determinant of autophagy synthesis (**Fig. 2E**). Moreover, this data suggests that amino acid levels must drop significantly from the amount in FM before there is an appreciable upregulation in autophagy.

**Figure 2.**
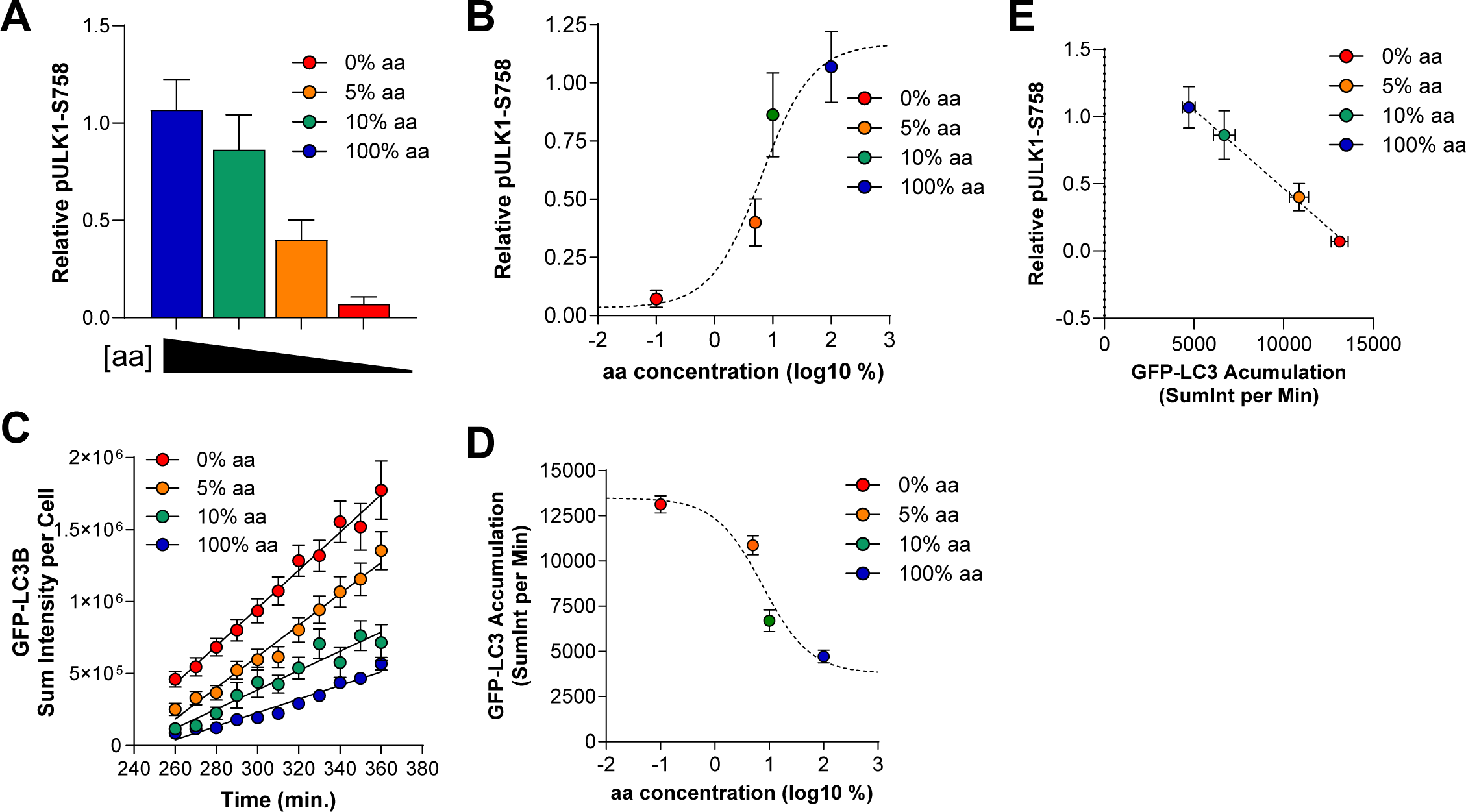
ULK1 phosphorylation and autophagy levels are tightly associated and regulated by amino acid levels. (**A-B**) U2OS cells were treated for 6 hours with FM (blue), indicated as 100% aa (the concentration found in RPMI-1640), or 10% (green), 5% (orange), or 0% (red) of that amino acid concentration. Cells were lysated and ULK1 phosphorylated at S758 quantified (relative to actin loading control and normalized to time 0 controls) (A). Bars represent means of 3 biological replicates. The data in (A) was fit to a sigmoidal dose-response curve (dashed line) to generate an EC_50_ of 6% aa (B). (**C-D**) Cells were treated with the medias described in A and imaged live from hours 4-6 in the presence of BafA1 (as in Fig. 1K). GFP-LC3 puncta were quantified from cells and sum intensity plotted (this is the sum of the intensity of all GFP-positive pixels, an output used to avoid potential issues with aggregated vesicles). Trajectories include mean objects per cell (symbols); bars represent s.e.m.; back lines represent simple linear regression (C). The data in (C) was fit to a sigmoidal dose-response curve (dashed line) to generate an EC_50_ of 7% aa (D). (**E**) The rate of BafA1-induced GFP-LC3 accumulation (derived from linear regression analysis, shown in (C) and the relative level of pULK1-S758 (from A) plotted to show a negative, linear association (dashed line, r^2^ = 0.815).

### PI(3)P effectors are differentially regulated during starvation

Upon aa starvation, cells immediately increased the abundance of puncta marked by WIPI1, a PI(3)P effector, increasing nearly 8-fold within the first hour (**Fig. 3A**). To determine whether these WIPI1-positive structures reflected newly synthesized PI(3)P-positive vesicles, we quantified puncta positive for EGFP-2xFYVE, a universal marker of PI(3)P-positive membranes in cells. We found that 2xFYVE-positive puncta were abundant in cells prior to starvation, with an average of 100 puncta per cell (**Fig. 3B**). During the first hour of aa-starvation, this abundance did not change, although it increased modestly later in starvation (**Fig. 3B**). This suggests that when cells are first aa-starved, cells respond by recruiting WIPI1 to pre-existing PI(3)P-positive membranes, potentially of endosomal origin as these are typically the most abundant EGFP-2xFYVE-positive structures^35^. In contrast to WIPI1, WIPI2B responded more slowly with robust increases in WIPI2B-positive puncta later in the aa starvation (**Fig. 3A**). Specifically, over half of the cells analyzed did not show an increase in WIPI2B puncta until 2-3 hours of aa starvation (see **Supplemental Table 7**).

**Figure 3.**
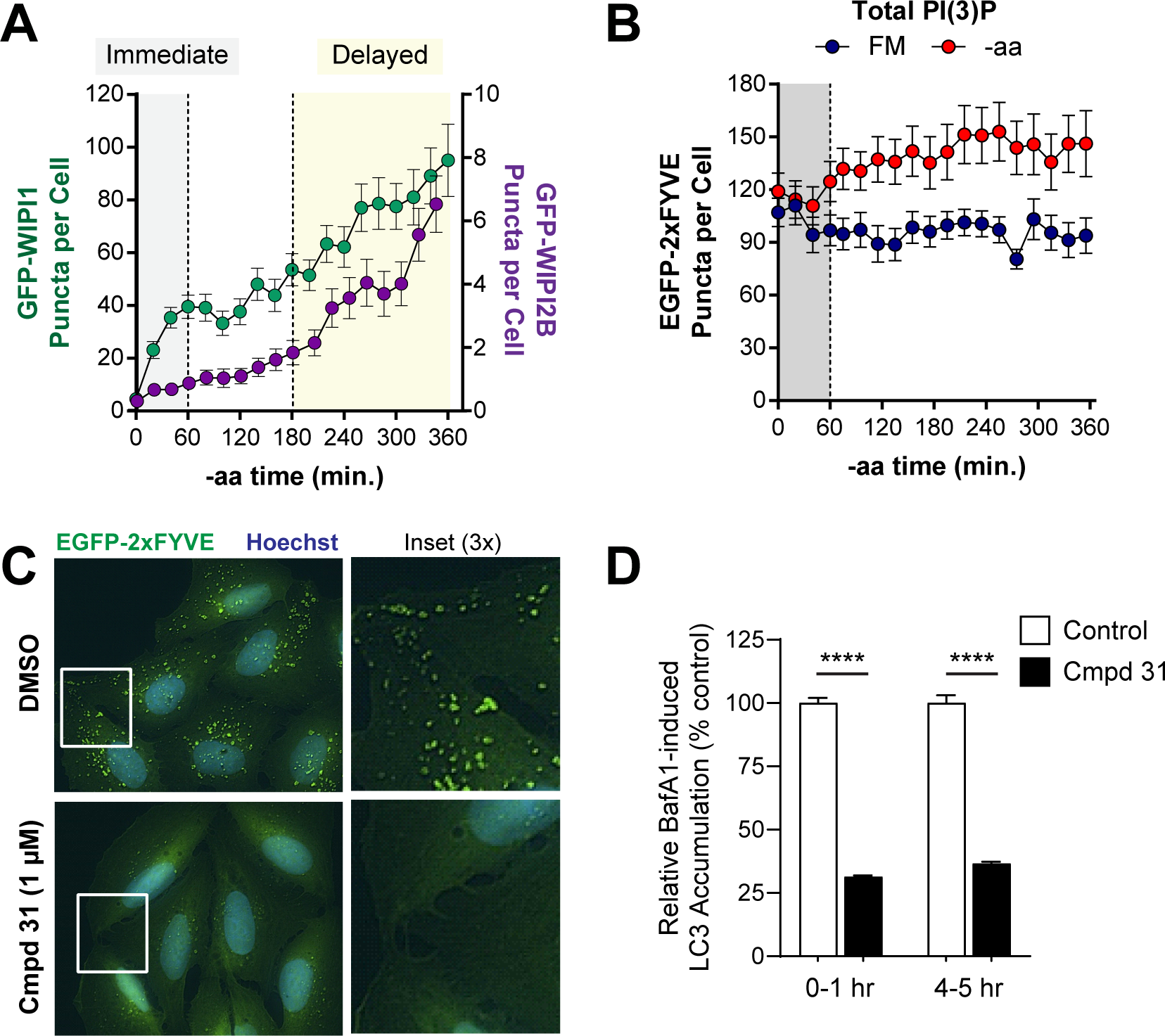
PI(3)P effectors, WIPI1 and WIPI2B, show distinct responses to aa starvation. (**A**) GFP-WIPI1 (green, left Y-axis) and GFP-WIPI2B (purple, right Y-axis) object counts over the 6 hour -aa treatment were overlaid. Gray region indicates immediate starvation period (0-1 hours) and yellow highlights period of delayed autophagy under sustained starvation (3-6 hours). (**B**) EGFP-2XFYVE puncta (PI(3)P-positive cell membranes) were quantified from cells under FM (blue) or -aa (red) treatment. Note a lack of substantial puncta increase in the immediate (0-1 hour) period (gray shading). (**C**) Representative EGFP-2xFYVE puncta in U2OS cells treated with a VPS34 inhibitor (1 μM compound 31, lower panels) or vehicle control (upper panels). Blue = Hoechst nuclear stain; green = EGFP-2XFYVE; captured with a 60x oil objective. (**D**) GFP-LC3 accumulation with BafA1 in the presence of compound 31 (1 μM) or vehicle control. BafA1 was added for 1 hour during either the first hour of -aa starvation (“0-1 hr” bars) or after 4 hours of -aa starvation (“4-5 hr” bars). Data shown represent GFP-LC3 puncta accumulating with the 1 hr BafA1 treatment relative to vehicle control. Symbols represent mean and bars are s.e.m. **** = adjusted p < 0.0001, one-way ANOVA.

We next wanted to establish the requirement for PI(3)P both early and late in aa-starvation (when WIPI1 and WIPI2B respond most dramatically, respectively). For this, we treated cells with a small molecule inhibitor of VPS34 (compound 31), the lipid kinase that produces PI(3)P. Compound 31 significantly reduced PI(3)P, as evidenced by loss of EGFP-2xFYVE puncta (**Fig. 3C**). Moreover, we observed an accompanying reduction in LC3-positive vesicle synthesis when measured both early (1 hour into aa-starvation) or late (5 hours into aa-starvation), consistent with these PI(3)P dynamics being critical for autophagy upstream of LC3 (**Fig. 3D**).

### Delayed autophagy is associated with transcriptional upregulation of machinery

We were intrigued by the observation that autophagy was particularly high late in starvation, beginning 3 to 4 hours into aa-deprivation, as evidenced by the increased abundance of nearly all markers measured (Fig. 1). Though amino acids were removed from the extracellular media at time 0, we wondered if the intracellular level of amino acids might shed light on the magnitude of autophagy, in particular, those known to regulate autophagy, including leucine, arginine, methionine, asparagine, and glutamine [REFs]. To address this, we employed LCMS metabolomics to quantify amino acids from cells 0 min, 20 min, 1 hour, or 4 hours into the aa-starvation. We found that there was nearly complete depletion of these amino acids within 60 minutes of aa-starvation (**Fig. 4A**) and no significant change in any amino acid between the 1 hour and 4 hour timepoints that might explain a change in autophagy (**Fig. 4B**).

**Figure 4.**
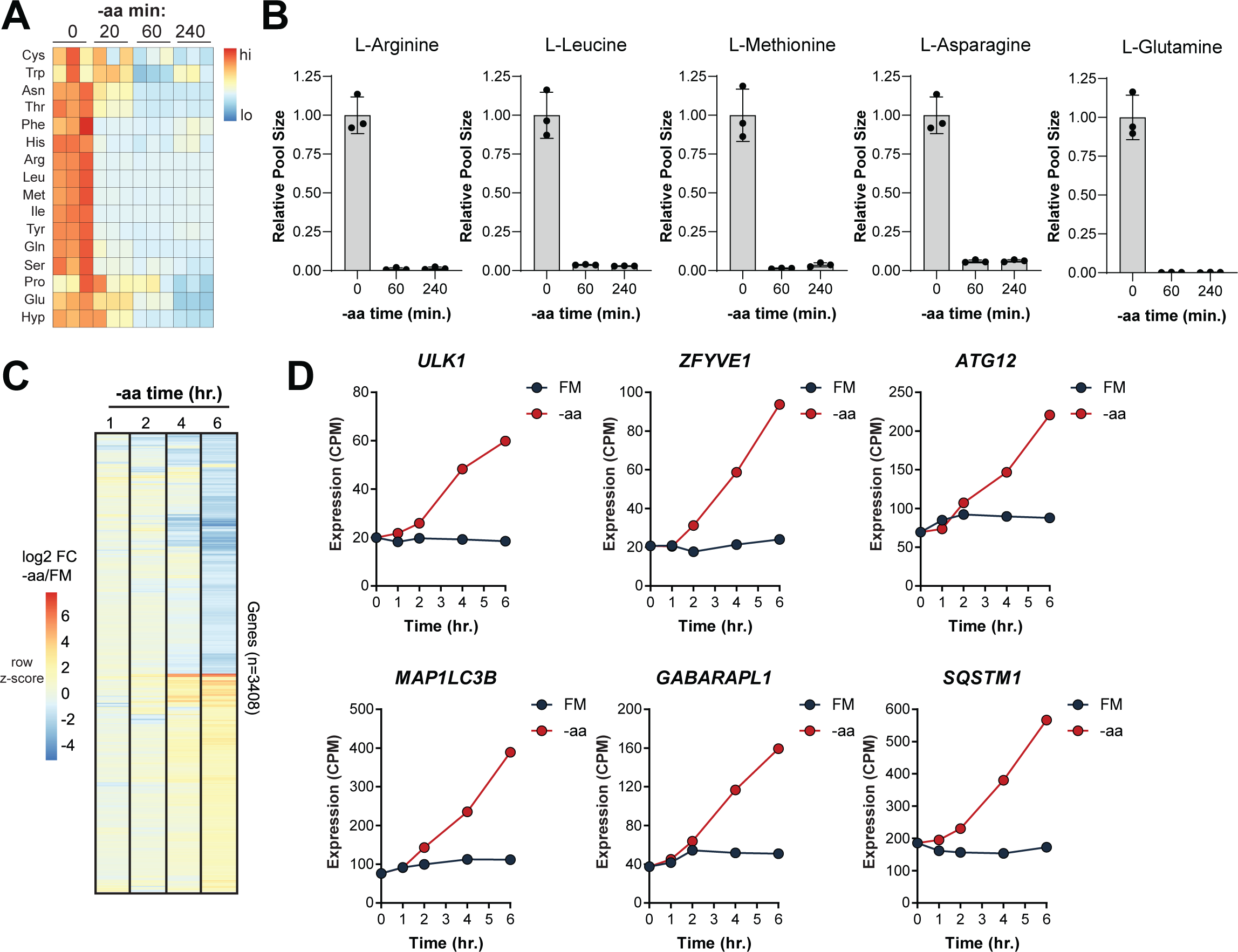
Autophagy machinery is transcriptionally upregulated with sustained amino acid starvation. (**A**) Intracellular amino acids were measured from triplicate samples after 0 min, 20 min, 1 hour, or 4 hours of -aa starvation using LCMS metabolomics. Relative levels (fold-change pool size relative to time 0) were calculated and displayed as a heatmap (red to blue, high to low; values are row-scaled z-scores). Note, we detected 16 of the 20 amino acids provided in FM. (**B**) Intracellular amino acids known to regulate MTORC1 and autophagy are plotted individually. Bars are mean (relative to pool size at time 0) and s.e.m. from triplicate samples (individual replicates shown as symbols). (**C**) RNAseq profiling was performed from cells cultured during -aa starvation for the indicate times compared to FM treatments. Differentially expressed genes (log2 fold-change +/-1 -aa versus FM at 6 hours) shown. (**D**) Expression of core autophagy genes (in counts per million, CPM) is shown at the indicated times of FM or -aa treatment.

Next, we reasoned that this timeframe may also be consistent with the induction of gene expression and transcription of autophagy genes^36,37^. To determine wether a transcriptional program might underly this increase in autophagy, we harvested RNA from cells after 1, 2, 4, or 6 hours of aa-starvation or after the same duration of treatment with FM. We then analyzed transcript expression by RNAseq. Indeed, we found a significant transcriptional reponse to aa-starvation (**Fig. 4C**) with 1,642 and 1,766 genes increased or decreased at least two-fold, respectively (**Supplemental Table 2**). Importantly, autophagy genes (identified in the Human Autophagy Database^38^) were significantly enriched among those increased (two-sided p = 0.0009, Fisher’s exact test). Specifically, we observed an upregulation of genes encoding core autophagy proteins, including *ULK1*, *ZFYVE1* (DFCP1), *ATG12*, *MAP1LC3B* (LC3B), *GABARAPL1* (an LC3-like molecule), and *SQSTM1* (p62, a selective autophagy cargo adaptor) (**Fig 4D**).

### Nutrient replenishment immediately restores cells to their basal autophagy level

After characterizing the response of cells to aa withdrawal, we wondered how quickly they would respond to the replenishment of amino acids following starvation. To test this, we starved cells of aa for 6 hours and then restored amino acid levels fully and quantified DFCP1, WIPI1, WIPI2B, or LC3 (in the presence of BafA1) for 1 hour (**Fig. 5A**). We found that all cells responded remarkably quickly to aa replenishment, with restoration of the basal state within 15-20 minutes (**Fig. 5B-G**). This suggests that cells are more buffered to the induction of autophagy in the face of starvation, showing a variable and often later autophagy induction, whereas they are primed to return to their basal homestatic rate of autophagy the moment a stressor, in this case aa-starvation, is relieved.

**Figure 5.**
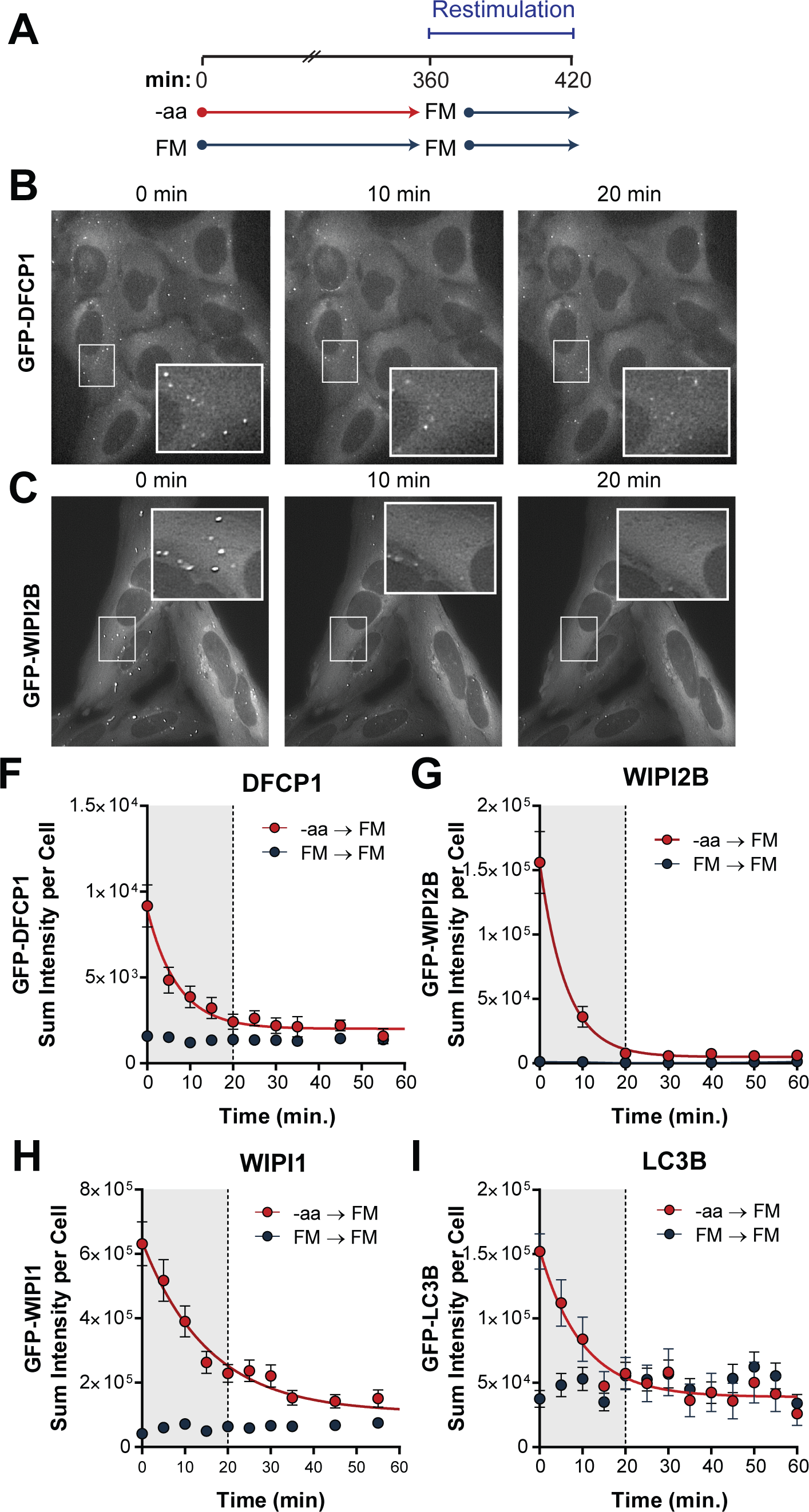
Autophagy levels are restored immediately upon amino acid replenishment. (**A**) Cells were cultured with or without amino acids for 6 hours prior to a restimulation phase of 60 min with FM (containing amino acids). (**B-C**) Representative images of GFP-DFCP1 (B) or GFP-WIPI2B (C) puncta in U2OS cells that were starved of amino acids for 6 hours and subject to aa-restimulation for 0 min (left), 10 min (middle) or 20 min (right). Insets show 2x magnification of indicates region to highlight disappearance of puncta. (**D-G**) DFCP1 (D), WIPI2B (E), WIPI1 (F), and LC3B (G) quantified from cells during the restimulation period following -aa (red) or FM (blue) treatments. Symbols are mean and bars are s.e.m. Solid lines are non-linear regression models (one phase exponential decay). All data presented as sum intensity (the sum of intensities of all GFP-positive pixels) to avoid any potential issues with aggregated vesicles. Gray shaded area emphasizes restoration to FM levels within 20 min of aa restimulation.

## DISCUSSION

Here, we have provided a detailed and quantitative description of autophagy induced by amino acid deprivation in a widely-used cancer cell line for signal transduction and autophagy research, including three protein phosphorylation sites and five fluorescent reporters for autophagy proteins. We supplemented these data with measurements of intracellular amino acids and mRNA transcript abundances. Collectively, we provide these data as a resource for the research community, for instance, to use in constructing or testing of computational models or for exploring single cell behavior across a population.

Our most notable observation was that U2OS cells strongly induce autophagy in response to loss of amino acids, but fail to do so in response to glucose starvation. In fact, we found slightly reduced LC3 vesicle dynamics during glucose starvation (see Fig. 1I-K). While contradictory to the original dogma that glucose starvation activates AMPK to upregulate autophagy, our data is consistent with recent data that suggest a more complex relationship between AMPK and autophagy^32^. By using carefully defined medias, we established an EC_50_ of approximately 6-7% aa towards pULK1-S758 and autophagy during a 6 hour cell culture treatment. This demonstrates that the amino acid concentrations found in typical cell culture media (like RPMI-1640, the basis for our medias) are in far excess of a threshold that would induce autophagy, consistent with the original intent of these medias to support cell viability and proliferation for several days in culture^39^.

While we draw conclusions from the population averages of data, we captured autophagy measurements from individual cells using live-cell imaging and analysis. Through this exercise, we uncovered significant cellular heterogeneity, despite each line being derived from a single clone. It is possible that the varied cellular responses we saw may relate to the cell cycle; for example, autophagy is inhibited during mitosis in order to protect nuclear contents during cell division, or they may relate to individual gene and protein expression in single cells^40^. A limitation of our study was that all markers were studied separately in individual cell lines, so it would be informative to express multiple effectors in a single cell line and confirm whether conserved subpopulations of cells exist that display distinct autophagy responses. Finally, it is intriguing to consider that individual cells responding at the level of autophagy on distinct timescales and with varied magnitudes could be beneficial to a cell population as a whole. While autophagy serves as an important cytoprotective mechanism in response to stress, it can also be detrimental if engaged too robustly or for a prolonged period of time; therefore, a diversity of autophagic responses may be favorable to a growing tumor.

We highlighted a clear difference in the reponse of two closely related PI(3)P effectors, WIPI1 and WIPI2B, to autophagy induction in our cells. It appears that WIPI1 is recruited immediately to abundant, pre-existing membrane structures while WIPI2B is induced later and with similar dynamics to DFCP1. Our data is consistent with WIPI2B localizing specifically to early autophagic membranes, and functioning closely with DFCP1^14^. In contrast, WIPI1 has been found to localize to not only early autophagic membranes, but also plasma membrane, nuclear membrane, endoplasmic reticulum (ER), and LAMP1-positive membranes^41^. We found that WIPI1-positive structures were far more abundant than WIPI2B-positive structures and closer to the number of total 2xFYVE-positive puncta, which labels all PI(3)P-positive membrane compartments, including the endolysosomal pathway. Thus, our data would support a model whereby WIPI1 is exquisitely sensitive to autophagy induction but functions at sites of existing and abundant PI(3)P, perhaps endosomal in origin, in contrast to WIPI2B, which may be more specific to the omegasome and is engaged most significantly after sustained starvation.

We observed an apparent boost in autophagy several hours into amino acid starvation, which we found co-occcurred with a transcriptional program that involved a diverse collection of differentially expressed genes. Among the most significantly upregulated genes were *ATF3*, *EGR1*, and *FOS*, which are immediate early genes known to be induced during amino acid deprivation as part of an amino acid response (AAR)^42^. In addition, of 198 autophagy genes detected in our cells, 36 (18%) were upregulated in contrast to just 7 (4%) downregulated. This supports a model whereby cells faced with persistant starvation increase autophagy machinery in order to maintain a high level of this process.

After hours of starvation, we found that cells responded swiftly and completely to the restoration of nutrients. In fact, all markers returned to basal levels within 10-20 minutes of amino acid replenishment, regardless of the initial level of autophagy in each cell. It is unclear if existing autophagic structures disassemble or proceed through completion, but regardless, this paints a picture of cells primed to return to their basal state of low autophagy. In this respect, autophagy can be viewed as a robust response to stress but one that is carefully regulated to avoid detrimental consequences.

## Supporting information

Supp Table 1 Formulations

Supp Table 2 RNAseq

Supp Table 3 DFCP1

Supp Table 3 WIPI1

Supp Table 5 WIPI2B

Supp Table 6 ATG5

Supp Table 7 LC3 FM BafA1

Supp Table 8 LC3 -aa BafA1

Supp Table 9 LC3 -Glc BafA1

Supp Table 10 LC3 FM NoBafA1

Supp Table 11 LC3 -aa NoBafA1

Supp Table 12 LC3 -Glc NoBafA1

Supp File 1 LCMS Details

## ACKNOWLEDGEMENTS

We thank current and former members of the MacKeigan laboratory for critical discussions and feedback. We thank the Van Andel Institute Cores for providing mass spectrometry, genomics, and bioinformatics facilities and services. We also thank William Hlavacek, Yen Ting, and Song Feng of the Los Alamos National Laboratory for their critical discussions and support for this work. This work was supported by grants and funding from the NIH to J.P.M. (R01CA197398). J.P.M. also has funding from the National Cancer Institute (R21CA270588, R21CA252430, and R21CA263133).

## METHODS

### Mammalian cell culture, reagents, and antibodies

The human osteosarcoma cell line U2OS (HTB-96) was purchased from American Type Culture Collection, and cells maintained in RPMI-1640 medium (Gibco, 11-875-119) supplemented with 10% fetal bovine serum (Corning, 35-010-CV) and cultured at 37°C in a humidified atmosphere containing 5% CO_2_. Cells were seeded 48 hours before the start of assays. We generated monoclonal U2OS cell lines expressing the following fluorescent plasmids: *1*) ptfLC3B was a gift from Tamotsu Yoshimori (Addgene plasmid #21074; http://n2t.net/addgene:21074; RRID:Addgene_21074^43^; *2*) pMXs-puro GFP-DFCP1 was a gift from Noboru Mizushima (Addgene plasmid #38269; http://n2t.net/addgene:38269; RRID:Addgene_38269^44^); *3)* pMXs-IP-EGFP-mATG5 was a gift from Noboru Mizushima (Addgene plasmid #38196; http://n2t.net/addgene:3819; RRID:Addgene_38196^45^). We obtained two additional monoclonal U2OS cell lines (GFP-WIPI1-U2OS and GFP-WIPI2b-U2OS) as a kind gift from Tassula Proikas-Cezanne^46^ and previously published U2OS-EGFP-2xFYVE^47^. Compound 31 (kind gift from Merck) and Bafilomycin A1 (BafA1; AG Scientific, B1183) stock solutions were prepared in dimethyl sulfoxide (DMSO) (Sigma-Aldrich, D2650). Equal concentrations of DMSO used for control treatments. Primary antibodies used were Total Acetyl-CoA Carboxylase (Cell Signaling Technology, 3662), Acetyl-CoA Carboxylase phospho-S79 (Cell Signaling Technology, 11818), S6K1 phospho-T389 (Cell Signaling Technology, 9205), ULK1 phospho-S758 (Cell Signaling Technology, 14202), and β-actin (Cell Signaling Technology, 3700). IRDye infrared fluorescent 680RD secondary goat anti-rabbit (926-68071) and anti-mouse (926-68070) purchased from LI-COR.

### Defined Medias

To prepare defined starvation medias, we reconstituted media containing the components of RPMI-1640, including the following reagents: 1XDPBS (Gibco, 14040-133), Phenol Red (Sigma-Aldrich, P3532), HEPES Buffer 1 M (Gibco, 15630-080), RPMI-1640 Vitamin Mix 100X (Sigma-Aldrich, R7256), Sodium bicarbonate (Sigma-Aldrich, S5761), reduced Glutathione (Sigma-Aldrich, G6013), 200 mM Glutamine (Gibco, 25030-081), D-Glucose (Fisher Scientific, D16), and dialyzed Fetal Bovine Serum (Sigma-Aldrich, F0392). Amino acid starvation was prepared by adding all components except amino acid mix and glucose starvation media was prepared by adding all components except glucose. See Supplemental Table 1 for full formulations. Medias were prepared, pH adjusted to 7.2-7.4, sterile-filtered, and stored at 4°C.

### Immunoblot Analysis

Cell lysates were prepared in ice-cold lysis buffer [40mM HEPES pH=7.4, 1 mM ethylenediaminetetraaceticacid (EDTA), 120 mM sodium chloride, 50 mM bis-glycerophosphate, 1.0% Triton-X 100, 1.5 mM sodium orthovanadate, 50 mM sodium fluoride, and protease inhibitor cocktail (Sigma-Aldrich, P8340)]. Lysate samples were clarified by centrifugation for 10 min at 13,400 rpm and 4°C. Total protein concentration was determined using Protein Assay Dye Reagent Concentrate (Bio-Rad, 5000006). Proteins resolved by pre-cast 4-12% NuPage Bis-Tris Plus Midi gels (Invitrogen, WG1403BOX) or 3-8% NuPage Tris-Acetate Midi gels (Invitrogen, WG1603BOX) and electrotransferred onto either nitrocellulose or polyvinylidene difluoride membranes. Membranes blocked with StartingBlock (TBS) Blocking Buffer (Thermo, 37542) and primary antibodies diluted in StartingBlock T20 (TBS) Blocking Buffer (Thermo, 37543) at 4°C. After three 5 min 0.1% Tween20-TBS washes, membranes incubated in secondary antibodies StartingBlock T20 (TBS) Blocking Buffer for 1 hour at room temperature. Protein bands imaged using LI-COR Odyssey Infrared Imaging System and quantified with LI-COR Image Studio Software.

### Fluorescence microscopy and vesicle quantification

Fluorescent cells (2.5 x 10^4^ cells per chamber) were seeded in 4-chamber 35mm CELLview dishes with glass bottom (Greiner Bio-One, 627870) and allowed to settle for 48 h before imaging. Thirty hours post-seeding, media in each chamber was replaced with fresh RPMI-1640 with 10%FBS. Forty-eight hours later cells were washed with warmed 1X DPBS (Gibco, 14190144) and given indicated media. In Figure 3, 1 µM of Compound 31 or vehicle control (DMSO) was administered at the time media was changed with amino acid starvation. For imaging, media was supplemented with 100 nM bafilomycin A1 (BafA1; A.G. Scientific) or an equivalent amount of vehicle (DMSO) for U2OS-ptfLC3B cell lines only for 2 hours as depicted in Figure 1H. All other cell lines contained only relevant starvation medias. For restimulation experiments, after media change cells were cultured for 6 hours in their respective starvation media, at the end of 6 hours imaging was paused to change all media to complete nutrient media. After all conditions had complete media imaging was resumed imaging every position every 5 minutes for 55 minutes. Cells were imaged live by maintaining a humid environment at 37°C and 5% CO_2_ in an environmental chamber fixed around the microscope stage. Five fields of view per chamber were chosen and NIS Elements software (Nikon) set to automatically image each position every 10 min for 2 hours (U2OS-ptfLC3B) or every 20 min for 6 hours (DFCP1, WIPI1, WIPI2b, and ATG5) using perfect focus to maintain the desired focal plane. One image for each point was obtained before the media change and defined as time zero. For restimulation experiments, media was replaced with complete nutrient media after 6 hours of starvation and imaging continued every 5 minutes for 55 minutes. Fields of view were chosen for their inclusion of healthy cells which were adherent and at the periphery of a cluster. Cells were imaged using a 60X oil objective, in the FITC channel, on a Nikon Ti Eclipse fluorescent microscope. Images were segmented into individual cells by defining regions of interest (ROIs). All images were deconvolved, top-hat transformed (peak identification), and thresholded (intensity) using NIS Elements Software to quantify EGFP-LC3-positive, GFP-DFCP1-positive, GFP-WIPI1-positive, GFP-WIPI2b-positive, EGFP-ATG5-positive, or EGFP-2xFYVE-positive puncta per cell^48^. Processed, quality data was obtained from an average of 58 cells per marker per media condition. Single cell data are provided in **Supplemental Tables 3-12**. Note, we confirmed LC3 puncta accumulation was specific to BafA1 by also collecting LC3 data under each condition with a vehicle (DMSO) control in place of BafA1. Though these data were not presented in the figures, the data is provided as Supp Tables 10-12.

### Amino Acid Measurements

180,000 U2OS cells were seeded per well of 6-well dishes in 2 ml RPMI-1640 + 10% FBS for 24 hours before being washed and incubated with with -aa media for 20 min, 1 hour, or 4 hours (a time 0 control was also harvested for comparison of aa levels). Media was aspirated, cells rinsed twice with ice-cold 0.9% sodium chloride (Sigma S8776), dried completely, and snap-frozen on dry ice. To extract intracellular polar metabolites, 1.5 mL of extraction solvent (40% acetonitrile, 40% methanol, 40% water, v/v) was added to frozen plates, plates were scraped, and extracts collected in 1.5mL Eppendorf tubes. Extracts were incubated on ice for 1 hour, then centrifuged at 15000xg for 10 min. 850µL of supertant was collected, dried in a vacuum evaporator, and resuspended in 50uL of water for LCMS analysis. An additional aliquot of 50µL from each supernatant was pooled, dried, and resuspended and anlyzed between every six experimental replicates to monitor instrument and sample drift during analysis. Metabolomics was performed using anion-paired chromatography on an Agilent 1290 UHPLC coupled to an Agilent 6470 QQQ mass spectrometer operated in ESI-in dMRM mode as described previously (PMID: 37095747, 36848144, 35981545, 31747582). Peak picking and integration was conducted in MassHunter (v8.0, Agilent). Complete instrument parameters, compound transition list, and peak integration notes are available in **Suppementary File 1**. Pool sizes of the indicated amino acids were normalized to the abundance in time 0 (non-starved) samples and plotted. All data was captured from 3 biological replicates per condition.

### RNAseq

U2OS cells were plated on 6-well dishes at 180,000 cells per well in 2 ml full media (FM) and incubated at 37°C for 24 hours. The next day, the plating media was aspirated, washed, and replaced with either FM or amino acid-starvation media. Cells were incubated for 1, 2, 4, or 6 hours at 37°C before cells lysed and RNA harvested using Qiagen QIAShredders and RNeasy spin columns following manufacturer’s protocols. RNA was quantified using a Qubit fluorimeter and analyzed for integrity using a Bioanalyzer. Total RNA was subject to polyA enrichment and sequenced with 2×50 bp reads on an Illumina NovaSeq (30M raw reads per library). For analysis, quality trimming and adapter removal was performed using Trim Galore, reads were mapped to the human reference genome using STAR, and quality control of trimming and alignment performed with MultiQC. We generated gene counts using STAR and imported into R and analyzed with an internal RNA-seq analysis pipeline (Van Andel Research Institute Bioinformatics and Biostatistics Core).

### Statistical Analysis

All statistical tests performed in GraphPad Prism. Biological replicates and error bars represent standard error of the mean (SEM), unless otherwise indicated. For Figure 4, Fisher’s exact test was performed by building a contingency table with 36 autophagy database genes and 16,06 non-autophagy database genes increased ≥1 log2 FC by 6 hours in -aa versus FM, and 162 autophagy database genes and 14,065 non-autophagy database genes not increased (two-tailed T-test, p = 0.0009). For Figure 3, one-way ANOVA performed with Sidak’s multiple comparison test; vehicle versus compound 31 compared at each timepoint.

## SUPPLEMENTAL INFORMATION

Supplemental Information includes 1 technical file and 12 tables and is available online.

## Supplemental File Legends

**Supplemental File 1**: LCMS instrument parameters, compound transition list, and peak integration notes for amino acid measurements.

**Supplemental Table 1**: Components and concentrations in FM, -aa, or -Glc medias.

**Supplemental Table 2**: RNAseq data from 1h, 2h, 4h, or 6h FM or -aa treatments. Values are log2-transformed counts per million (CPM). Fold change (FM vs -aa at 6hrs) is also included and presence or absence in the Human Autophagy Database.

**Supplemental Tables 3-6:** Each file holds 18 sheets; 6 sheets for each of 3 conditions. Each of the following sheet names is appended with “FM”, “-aa”., or “-Glc”: 1) *Information* (experimental notes), 2) *Number of objects* (puncta count per cell), 3) *ROI area* (area of each cell from which puncta were quantified), 3) *Sum Intensity* (sum of intensity of all pixels in puncta), 4) *Binary Area* (total area taken up by puncta in cell), 5) *Mean Intensity* (average fluorescent intensity of pixels in puncta). Individual cells are represented on rows.

**Supplemental Tables 7-12:** These files contain GFP-LC3 data collected during treatment with BafA1 (Supplemental Tables 7-9) or with a vehicle (DMSO) control (Supplemental Tables 10-12). There is a file for each condition (FM, -aa, or -Glc, noted in file title) and within each, 6 sheets described above are included for each of 3 timepoints (0-2 hr, 2-4 hr, 4-6 hr). The sheet names are appended with these times.

## Notes

### Competing Interest Statement

J.P.M. has consulting agreements with Merck and scholarly activity with the Translational Genomics Research Institute (a non-profit organization). R.G.J. is a scientific advisor for Agios Pharmaceuticals and Servier Pharmaceuticals and is a member of the Scientific Advisory Board of Immunomet Therapeutics.

